# Improving the computation efficiency of polygenic risk score modeling: Faster in Julia

**DOI:** 10.1101/2021.12.27.474263

**Authors:** Annika Faucon, Julian Samaroo, Tian Ge, Lea K. Davis, Ran Tao, Nancy J. Cox, Megan M. Shuey

## Abstract

To enable large-scale application of polygenic risk scores in a computationally efficient manner we translate a widely used polygenic risk score construction method, Polygenic Risk Score – Continuous Shrinkage (PRS-CS), to the Julia programing language, PRS.jl. On nine different traits with varying genetic architectures, we demonstrate that PRS.jl maintains accuracy of prediction while decreasing the average run time by 5.5x. Additional programmatic modifications improve usability and robustness. This freely available software substantially improves work flow and democratizes utilization of polygenic risk scores by lowering the computational burden of the PRS-CS method.

## INTRODUCTION

The conceptual framework known as the ‘liability-threshold model’ asserts that complex diseases have many contributing variants of small effect, which collectively contribute to a continuous distribution of genetic liability in a population. Thus, when a large enough collection of risk alleles are aggregated in an individual together with environmental risk factors such that they pass a critical threshold, the complex disease will manifest (Falconer 1965). The additive genetic portion of this liability attributable to common variants can be estimated with a polygenic risk score (PRS). A PRS is generally calculated as a weighted sum of risk alleles present in an individual genome, where the weights are defined by the effects estimated in genome wide association studies (GWASs).(Chatterjee et al. 2016)

Since the advent of PRS methods, various studies have proven their potential to improve health by informing therapeutic intervention,(Tikkanen et al. 2013; Mega et al. 2015) disease screening,(Hsu et al. 2015) and lifestyle choices for a multitude of polygenic conditions. In fact, polygenic risk scoring has long been at the center of genetic research (MultiBLUP (Speed and Balding 2014), PLINK (Purcell et al. 2007), PRSice (Choi and O’Reilly 2019), LDpred (Vilhjalmsson et al. 2015)). In simulation and real data analyses, PRS-CS was demonstrated as a top performing method (Ge et al. 2019; Pain et al. 2021). Despite their popularity and importance, PRS methods need development, particularly related to computational expense. As large datasets become publicly available and computation moves to the cloud (Langmead and Nellore 2018), research demands the use of computational programs that can scale and cost-effectively utilize resources. Because the Julia programming language has consistently demonstrated increased efficiency of computation over other programming languages, along with other advantages (Bezanson et al. 2018), we created a Julia translation of the commonly used Python based PRS-CS program, PRS.jl.

Below we introduce the PRS.jl program and benchmark it against PRS-CS, tracking model accuracy and computational improvements across nine well-characterized polygenic phenotypes including both continuous and binary outcomes.

## RESULTS

### PRS.jl performance overview

PRS.jl is a direct translation of the PRS-CS Python program into the Julia language. This translation improves the computational efficiency of PRS estimation across a variety of polygenic traits.

Using the auto global shrinkage calculation with 10,000 MCMC iterations on a single Haswell node, 8 CPUs available total, and a maximum memory allocation of 80 GB, we observed an average 5.5x improvement in computational speeds when using PRS.jl compared to PRS-CS across nine phenotypes. The improvements in speed ranged from 3.8x to 6.4x (Table 1). For the quantitative phenotypes - body mass index (BMI), high density lipid cholesterol, low-density lipid cholesterol, total cholesterol, triglycerides, and estimated glomerular filtration rate (eGFR) - the average improvement was 5.6x. For the binary traits - asthma, coronary artery disease, and type 2 diabetes mellitus - the improvement was 5.1x. These reported computational times represent the total amount of run time when chromosomes are sequentially analyzed; processing the chromosomes in parallel can substantially reduce time to results.

**Table 1.** Individual and average run times for PRS-CS and PRS.jl by phenotype.

|  | PRS-CS |  |  |  | PRS.jl |  |  |  | Average |
| --- | --- | --- | --- | --- | --- | --- | --- | --- | --- |
|  | Run 1 | Run 2 | Run 3 | Mean (SD) | Run 1 | Run 2 | Run 3 | Mean (SD) | Improvement* |
| <u>Quantitative Phenotypes</u> |  |  |  |  |  |  |  |  |  |
| Body Mass Index | 62:21:25 | 65:12:16 | 59:41:13 | 62:24:58 (1:28:32) | 12:01:15 | 12:20:31 | 11:19:32 | 11:53:46 (0:31:10) | 5.3 |
| Cholesterol | 56:51:43 | 55:36:35 | 59:12:59 | 57:13:46 (1:49:52) | 12:07:50 | 8:15:23 | 12:34:38 | 10:59:17 (2:22:34) | 5.2 |
| eGFR | 65:52:20 | 69:43:40 | 77:57:50 | 71:11:17 (6:10:36) | 13:54:28 | 15:57:40 | 14:01:35 | 14:37:54 (1:09:10) | 4.9 |
| High-density lipoprotein | 59:58:13 | 56:39:56 | 59:19:08 | 58:39:06 (1:45:02) | 8:21:45 | 8:16:13 | 10:58:48 | 9:12:15 (1:32:19) | 6.4 |
| Low-density lipoprotein | 63:38:06 | 56:17:24 | 56:47:47 | 58:54:26 (4:06:08) | 8:26:04 | 8:18:47 | 12:54:02 | 9:52:58 (2:36:51) | 6.0 |
| Triglycerides | 58:00:52 | 56:04:48 | 60:45:49 | 58:17:10 (2:21:13) | 8:18:01 | 8:09:24 | 11:48:25 | 9:25:17 (2:04:02) | 6.2 |
| <u>Binary Phenotypes</u> |  |  |  |  |  |  |  |  |  |
| Asthma | 41:58:26 | 40:15:14 | 41:29:02 | 41:14:14 (0:53:10) | 5:26:44 | 6:25:06 | 7:27:54 | 6:26:35 (1:00:36) | 6.4 |
| Coronary Artery Disease | 66:59:45 | 66:00:16 | 64:05:35 | 65:41:52 (1:28:32) | 10:43:53 | 15:05:15 | 13:04:09 | 12:57:46 (2:10:48) | 5.1 |
| Type 2 Diabetes Mellitus | 68:35:38 | 69:10:26 | 68:08:43 | 68:38:16 (0:30:57) | 17:03:18 | 20:13:24 | 16:38:33 | 17:58:25 (1:57:33) | 3.8 |
| <u>All Phenotypes Combined</u> |  |  |  |  |  |  |  |  |  |
|  | 544:16:28 | 535:00:35 | 547:28:06 | 542:15:03 (6:28:16) | 96:23:18 | 103:01:43 | 110:47:36 | 103:24:12 (7:12:35) | 5.5 |

All run times are presented as hour:minutes:seconds
\*Average improvement is estimated as the mean PRS-CS / mean PRS.jl

We next demonstrate that these improvements in speed did not come at the expense of PRS accuracy (Figure 1).

**Figure 1.**
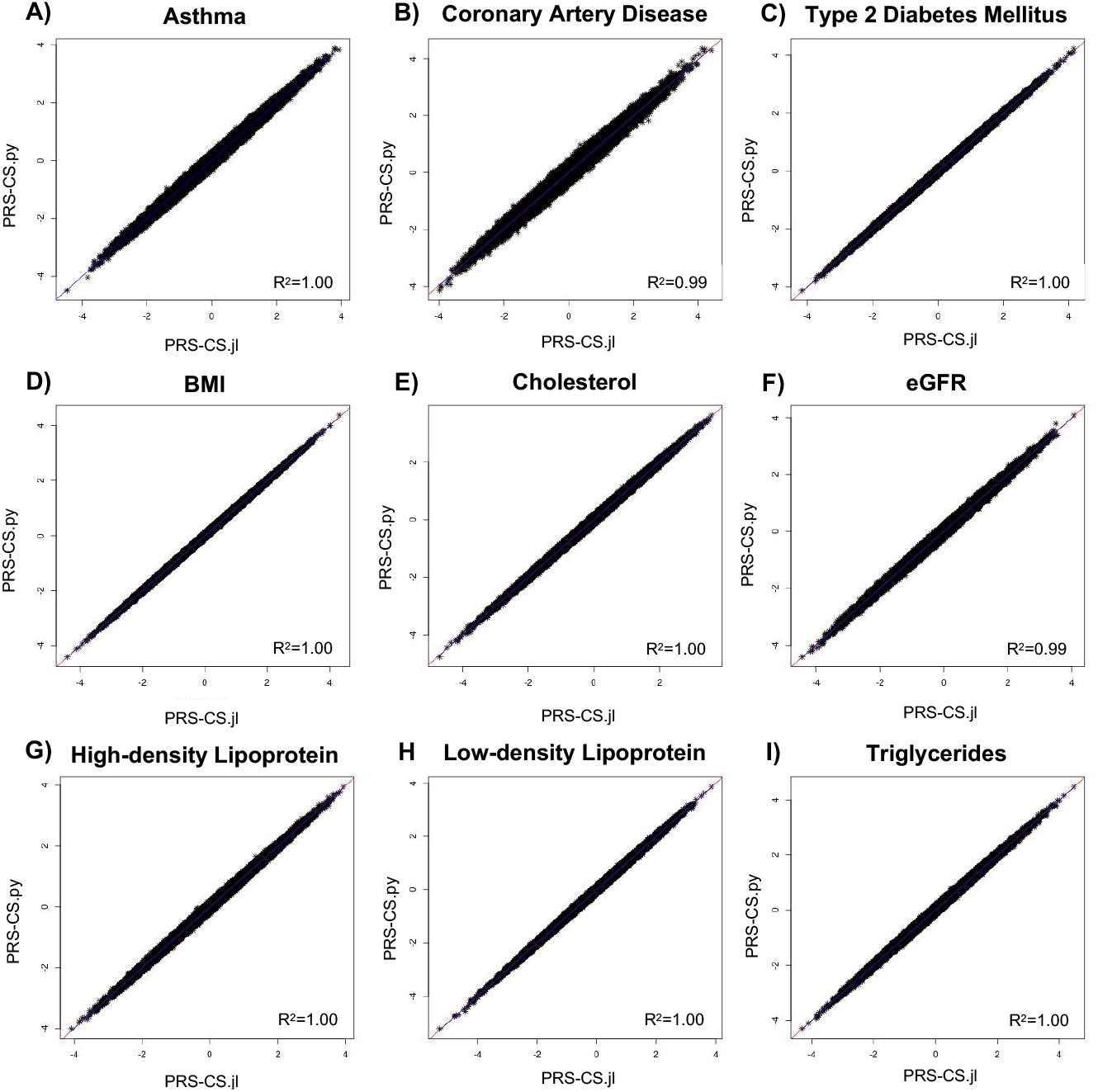
Plots comparing PRS-CS.py and PRS.jl polygenic risk score estimates for each trait. Plots of the polygenic risk scores calculated by PRS-CS on the y-axis compared to the scores calculated by PRS.jl on the x-axis for each trait: A) Asthma, B) Coronary Artery Disease, C) Type 2 Diabetes Mellitus, D) Body Mass Index (BMI), E) Cholesterol, F) estimated glomerular filtration rate (eGFR), G) High-density lipoprotein, H) Low-density Lipoprotein, and I) Triglycerides. The correlation R^2^ are presented in the corner of each plot.

Next, we show the retained accuracy of PRS.jl estimate for given phenotypes by demonstrating the consistency of posterior SNP weights and the resulting PRSs compared to PRS-CS. Specifically, to assess the consistency of SNP weights, we calculate the squared error for each SNP between the PRS.jl and PRS-CS output (Table 2). The median squared error between the two algorithms ranged from 2.00 E-11 and 6.83 E-11 across phenotypes, with a median of 3.00 E-11. This is similar to the median squared errors within the same program on different runs. A t-test comparing the posterior SNP effect sizes estimated by PRS.jl and PRS-CS found no statistically significant difference (all p>0.85).

**Table 2.**
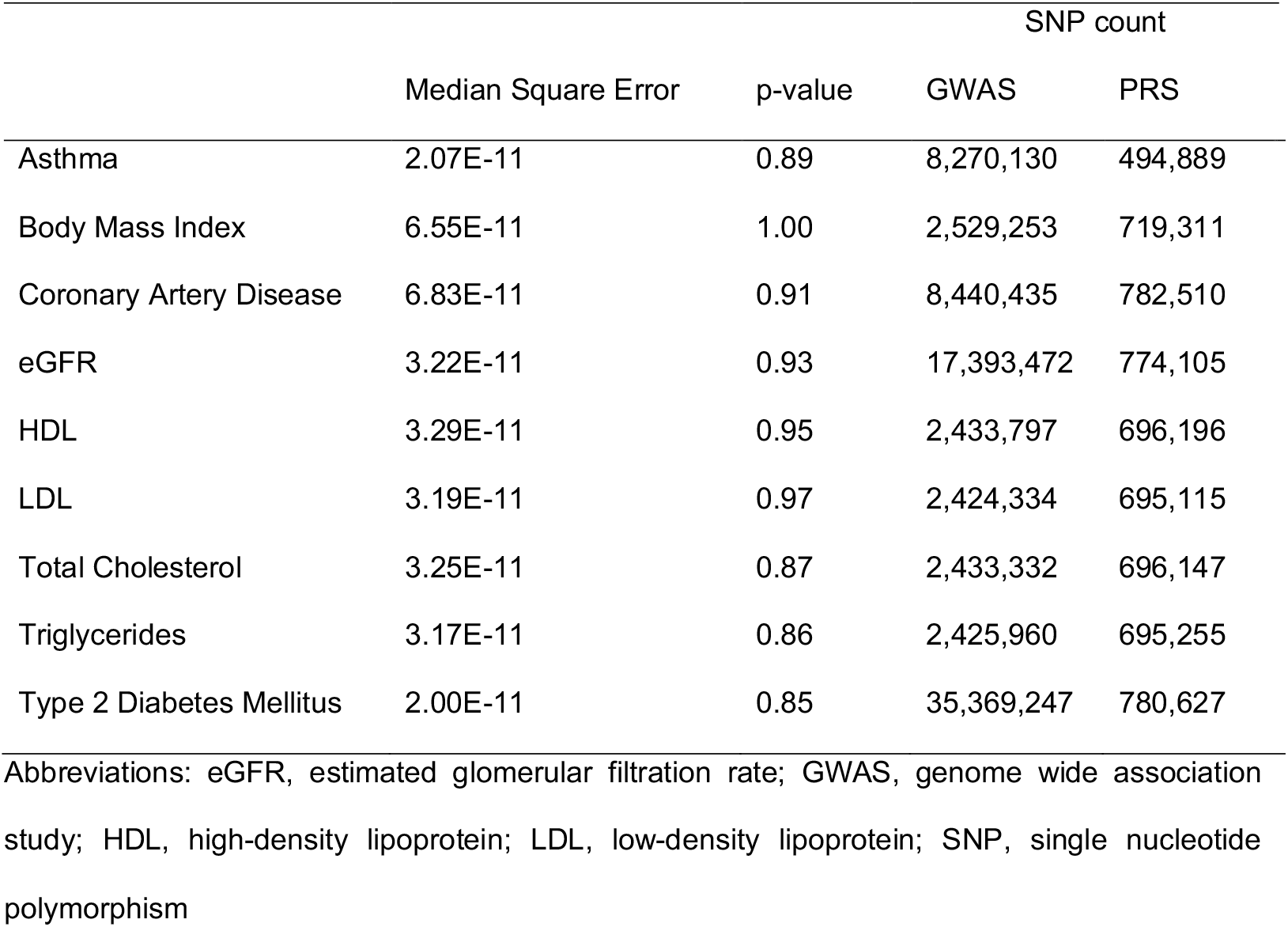
Median Squared Error and p-value from the T-test comparing SNP weights between python and Julia implementations for a single run.

|  | Median Square Error | p-value | SNP count |  |
| --- | --- | --- | --- | --- |
|  |  |  | GWAS | PRS |
| Asthma | 2.07E-11 | 0.89 | 8,270,130 | 494,889 |
| Body Mass Index | 6.55E-11 | 1.00 | 2,529,253 | 719,311 |
| Coronary Artery Disease | 6.83E-11 | 0.91 | 8,440,435 | 782,510 |
| eGFR | 3.22E-11 | 0.93 | 17,393,472 | 774,105 |
| HDL | 3.29E-11 | 0.95 | 2,433,797 | 696,196 |
| LDL | 3.19E-11 | 0.97 | 2,424,334 | 695,115 |
| Total Cholesterol | 3.25E-11 | 0.87 | 2,433,332 | 696,147 |
| Triglycerides | 3.17E-11 | 0.86 | 2,425,960 | 695,255 |
| Type 2 Diabetes Mellitus | 2.00E-11 | 0.85 | 35,369,247 | 780,627 |

Abbreviations: eGFR, estimated glomerular filtration rate; GWAS, genome wide association study; HDL, high-density lipoprotein; LDL, low-density lipoprotein; SNP, single nucleotide polymorphism

Lastly, we examined accuracy of the PRSs relative to the traits measured in BioVU. For the quantitative traits, we compared prediction accuracy by R^2^ between the observed and predicted phenotypes in the BioVU testing set (Table 3). For the binary traits, we compared prediction accuracy by Area Under the Curve (AUC), the Nagelkerke R^2^, and the odds ratio of the top 10% versus the remaining 90% (Table 4). PRS-CS and PRS.jl had nearly identical accuracies across all tested traits.

**Table 3.** Comparison of PRS-CS and PRS.jl performance for quantitative traits using, as covariates, age, sex, and PCs 1-10.

|  | R<sup>2</sup> |  | Number of subjects |
| --- | --- | --- | --- |
|  | PRS-CS | PRS.jl |  |
| Body Mass Index | 0.1141(<0.0001) | 0.1141 (<0.0001) | 60,584 |
| Cholesterol | 0.0657 (0.0002) | 0.0657 (0.0002) | 34,347 |
| eGFR | 0.4921 (<0.0001) | 0.4922 (<0.0001) | 34,797 |
| High-Density Lipoprotein | 0.1089 (<0.0001) | 0.1088 (<0.0001) | 33,338 |
| Low-Density Lipoprotein | 0.0835 (<0.0001) | 0.0835 (0.0001) | 32,061 |
| Triglycerides | 0.2326 (<0.0001) | 0.2326 (<0.0001) | 34,531 |

All data is presented as: mean (standard deviation)

**Table 4.** Comparison of PRS-CS and PRS.jl performance for binary traits.

|  | Nagelkerke R<sup>2</sup> |  | Area Under the Curve |  | 10% Odds Ratio |  | Number of subjects |  |
| --- | --- | --- | --- | --- | --- | --- | --- | --- |
|  | PRS-CS | PRS.jl | PRS-CS | PRS.jl | PRS-CS | PRS.jl | Cases | Controls |
| Asthma | 0.0176 (<0.0001) | 0.0176 (<0.0001) | 0.560 (<0.0001) | 0.560 (0.0002) | 1.54 (0.02) | 1.54 (0.02) | 8,210 | 64,618 |
| Coronary Artery Disease | 0.3212 (0.0001) | 0.3214 (0.0001) | 0.552 (0.0004) | 0.552 (0.0005) | 1.71 (0.02) | 1.73 (0.01) | 16,807 | 56,021 |
| Type 2 Diabetes Mellitus | 0.1715 (<0.0001) | 0.1716 (<0.0001) | 0.626 (0.0001) | 0.626 (0.0001) | 2.75 (0.017) | 2.77 (0.008) | 13,688 | 59,140 |

All data is presented as: mean (standard deviation)

## DISCUSSION

Polygenic risk scores are hailed for their potential to revolutionize clinical and precision medicine. Despite early successes there remains considerable concerns relating to the broader applicability of PRSs to genetically diverse populations as well as the computational power required to utilize these approaches at scale. With the growing availability of large-scale biobanks including: All of Us (All of Us Research Program et al. 2019), Biobank Japan (Nagai et al. 2017), FinnGen (Consortium 2021), and UKBiobank (Bycroft et al. 2018) the need for improved genomic analysis tools that have the potential to handle these larger sample sets in a faster, less computationally intensive manner without sacrificing efficacy is paramount. The Julia programming language has many features that allow for these improvements including an efficient type-system and multiple dispatch, a variety of optimized matrix routines, and a straightforward application programming interface (API) for accessing single instruction, multiple data (SIMD) and multi-threading. Here, we demonstrate how a basic port of the commonly used PRS-CS package to the Julia language, PRS.jl, can improve the program’s speed without sacrificing PRS estimation across a variety of traits. PRS.jl is freely available for download through GitHub (github.com/fauconab/PolygenicRiskScores.jl) and is a drop-in replacement for PRS-CS. The available README text instructs even novice Julia programming language users how to execute this software with ease. Because of the usability and performance improvements, we believe PRS.jl will allow for broader and more efficient use of PRSs in genomic medicine.

Further, the development of tools for genomic analyzes that are both fast and computationally efficient, such as PRS.jl, have the potential to democratize genomic research. To date, human genomics research is over-represented by high-income countries, which tend to have more powerful computational resources and greater funding for the sciences. The lack of population diversity and global representation is driven by many factors; however, a lack of resources, financial and human, are consistently noted as key limitations that prevent middle- and low-income countries from fully utilizing and contributing to genomic research (Marques-de-Faria et al. 2004; Hardy et al. 2008; Seguin et al. 2008; Kaur et al. 2019). Due to this disparity, these countries could benefit the most from advances in genomic medicine.

Knowing the impact, utility, and potential of PRSs to drive personalized medicine while acknowledging the immense bias in data availability and utilization, the National Institutes of Health funded a large initiative to fund PRS research in diverse populations. By reducing the computational needs of key algorithms, low resourced research groups can utilize these algorithms to benefit their scientific endeavors and provide potential benefit for their populations. The current versions of PRS-CS and PRS.jl are limited in their utility in ancestrally diverse populations (Duncan et al. 2019), a limitation that has been addressed by PRS-CSx (Huang et al. 2021). Future work to extend Julia improvements to the PRS-CSx framework for faster trans-ancestry PRS calculations has been planned.

The PRSs in this set of work are derived using standard inputs and publicly available summary statistics. Different discovery GWASs for the same traits can provide different polygenic risk estimates at the individual level (Schultz et al. 2021); therefore, association of scores to clinical values in this paper may be different than other papers using similar clinical traits due to differences in the discovery summary statistics. As such, the PRSs generated in this paper do not represent the most optimized PRS for any particular trait. Despite this limitation in study design, our work clearly demonstrates that the accuracy for both the Python and Julia versions of PRS-CS are nearly identical. Because the base datasets and testing populations are identical, our design allows for a head-to-head comparison of PRS-CS versus PRS.jl. Additional study limitations include the utilization of a direct translation of the PRS-CS package. While this approach allows us to directly compare the accuracy performance across the Python and Julia implementations, it does not fully utilize the various computational improvements that Julia affords. For example, future work aims to utilize Julia’s multi-threading and GPU compute capabilities, are effective methods for computational acceleration of programs which heavily utilize matrix operations. These additional compute capabilities would allow PRS.jl to better utilize the hardware that users have available, and make processing of even larger data sets feasible.

## METHODS

### PRS.jl development

PRS-CS was cloned from https://github.com/getian107/PRScs. This implementation was translated to the Julia programming language. Development of PRS.jl was carried out in the open, with all contributions being publicly posted to the PRS.jl GitHub repository.

### Training dataset and example phenotypes

We used the Vanderbilt University Medical Center Synthetic Derivative (VUMC SD), a deidentified copy of the electronic health record (EHR), for the identification of the nine test phenotypes. VUMC is as a tertiary care center that provides inpatient and outpatient care in Nashville, TN. The SD includes more than 2.8 million patient records, that contain International Classification of Diseases, 9^th^ and 10^th^ editions (ICD9 and ICD10), codes; Current Procedural Terminology (CPT) codes; laboratory values; medication usage; and clinical documentation (Roden et al. 2008). From the SD a subset of patients are part of VUMC BioVU, a biobank that links the de-identified EHRs of patients to discarded blood samples for the extraction of genetic materials (Roden et al. 2008). The VUMC Institutional Review Board oversees BioVU and approved these projects.

#### Genotyping and Quality Control

We obtained genome-wide data from 94,474 BioVU individuals genotyped on the Illumina MEGA^EX^ array. We used PLINK v1.9 to filter genotypes with low SNP (<0.95) call rate, and individuals with low call rate (<0.98), sex discrepancies, and excessive heterozygosity (|Fhet|>0.2). Principal component analysis on the genotyped BioVU cohort together with individuals from the 1000 Genomes Project (Genomes Project et al. 2015) were used to create CEU-YRI and CEU-CHB axes in FlashPCA2. Simple thresholding was used to select individuals of recent European ancestry. We confirmed the absence of genotyping batch effects through logistic regression with ‘batch’ as the phenotype. We used the Michigan Imputation Server (Das et al. 2016) with the reference panel from the Haplotype Reference Consortium to impute genotypes. SNPs were filtered for imputation quality (R^2^ > 0.3 or INFO > 0.95) and converted to hard calls. We restricted PRS calculations to autosomal SNPs with minor allele frequency (MAF) above 0.01. We removed SNPs that differed by more than 10% in MAF from the 1000 Genomes Project phase 3 CEU (Genomes Project et al. 2015) set and those with a Hardy Weinberg Equilibrium p<10^−10^. The resulting data set contained hard-called SNP information for 9,386,383 SNPs in 72,828 individuals of European Ancestry.

### Polygenic risk score calculations

We calculate PRSs for individuals using PRS-CS (Ge et al. 2019) and our translation of the package to the Julia programming language, PRS.jl. PRS-CS/PRS.jl uses Bayesian regression with a continuous shrinkage prior to model polygenic effects on the phenotype and updates the weight of each SNP within each LD block in posterior inference. The program can use an assigned global shrinkage parameter, or automatically learn the parameter from the data.

#### Model Performance in BioVU

Summary statistics were downloaded for six quantitative traits: BMI, high-density lipid cholesterol, low-density lipid cholesterol, total cholesterol, triglycerides, and eGFR (Willer et al. 2013; Hellwege et al. 2019; Pulit et al. 2019) and three binary traits: asthma, coronary artery disease, and type 2 diabetes Mellitus (T2DM) (Preuss et al. 2010; Zhu et al. 2019; Vujkovic et al. 2020).

Summary statistics were processed to get these input files in a format that the original PRS-CS method can accept (columns reordered and renamed using R). Polygenic risk scores were calculated in triplicate using a single CPU architecture. The scripts used to call both programs are available at https://juliahub.com/ui/Packages/PolygenicRiskScores/zm2vm/0.1.0.

### Computational performance comparison

All computations were performed using Vanderbilt University’s Advanced Computing Center for Research and Education (ACCRE, www.accre.vanderbilt.edu). Each PRS run was restricted to a single Haswell node with an allocation of 8 CPUs and 80 GB of memory. To minimize run-time variabilities related to cluster usage we initiated the PRS-CS and PRS.jl runs for each phenotype simultaneously. Subsequent benchmarking runs were initiated over a three-month time course. The processing time for each PRS run was recorded. The mean and standard deviation of the three runs per phenotype were calculated.

### PRS.jl and PRS.py Performance Comparison

Sensitivity analyses demonstrated similar output for the two methods when using a fixed global shrinkage parameter or the auto algorithm. Thus, in each case, we used the auto version, allowing for the estimation of the global shrinkage parameter from the data. Once the posterior beta values were calculated, PLINK v1.9 was used to score each individual. PRSs for each phenotype were scaled to have mean zero and unit standard deviation using the built-in R scale() function. Prediction accuracy was assessed using real phenotypic values in BioVU and covariate adjustment (sex, age, and PCs 1-10).

#### Verification of Quantitative trait Performance in BioVU

Accuracy of PRSs calculated from quantitative trait summary statistics was assessed using R^2^ between the scaled PRSs and median values by person from BioVU data processed by a previously published quality control pipeline, called Quality Lab (Dennis et al. 2021).

#### Verification of Binary trait Performance in BioVU

Accuracy of PRSs trained from binary trait summary statistics was assessed using the AUC, the Nagelkerke R^2^, and the odds ratio of the top-10% compared against the bottom 90% between the scaled PRSs and the binary presence of clinical codes that are representative of the clinical disease. The specific codes used for asthma and coronary artery disease are available in Supplemental Table 1. These codes include ICD-9 and -10 codes that mirror the clinical disease. For type 2 diabetes mellitus, however, presence of the condition was determined using an updated version of a previously published phenotyping algorithm which is effective at distinguishing type 1 and type 2 diabetes (Pacheco and Thompson 2012).

## DATA ACCESS

GWAS summary statistics were downloaded from publicly available resources. BioVU summary statistics are made available upon reasonable request to authors. PRS-CS/PRS.jl is available for download from GitHub (https://github.com/fauconab/PolygenicRiskScores.jl).

## COMPETING INTEREST STATEMENT

Authors declare no competing interests.

## ACKNOWLEDGMENTS

The work was supported by the following grants: 2R01CA157823-07A1 and R01HL151152. The dataset used for performance characterization was obtained from the Vanderbilt University Medical Center Synthetic Derivative, which is supported by institutional funding, the 1S10RR025141-01 instrumentation award, and by the CTSA grant UL1TR000445 from National Center for Advancing Translational Sciences/National Institutes of Health.

## AUTHOR CONTRIBUTIONS

AF and JS conceived the project. LKD, NJC, and MMS provided critical oversight of project execution and materials. AF, JS, LKD, NJC, and MMS were involved in data analysis, interpretation of results, and manuscript preparation. All authors were involved in project development, critical manuscript review, and approval.

## Notes

### Competing Interest Statement

The authors have declared no competing interest.

